# Altered somatic hypermutation patterns in COVID-19 patients classifies disease severity

**DOI:** 10.1101/2022.12.20.521139

**Authors:** Modi Safra, Zvi Tamari, Pazit Polak, Shachaf Shiber, Moshe Matan, Hani Karameh, Yigal Helviz, Adva Levy-Barda, Vered Yahalom, Avi Peretz, Eli Ben-Chetrit, Baruch Brenner, Tamir Tuller, Meital Gal-Tanamy, Gur Yaari

**Author notes:** Correspondence: Gur Yaari.

## Abstract

The success of the human body in fighting SARS-CoV-2 infection relies on lymphocytes and their antigen receptors. Identifying and characterizing clinically relevant receptors is of utmost importance. We report here the application of a machine learning approach, utilizing B cell receptor repertoire sequencing data from severely and mildly infected individuals with SARS-CoV-2 compared with uninfected controls. In contrast to previous studies, our approach successfully stratifies non-infected from infected individuals, as well as disease level of severity. The features that drive this classification are based on somatic hypermutation patterns, and point to alterations in the somatic hypermutation process in COVID-19 patients. These features may be used to build and adapt therapeutic strategies to COVID-19, in particular to quantitatively assess potential diagnostic and therapeutic antibodies. These results constitute a proof of concept for future epidemiological challenges.

## Background

Despite the unprecedented speed of vaccine development against SARS-CoV2, the virus continues to undergo changes that cause repeated waves of COVID-19 morbidity worldwide, with increasing infectivity. Risk factors such as age (> 60) and preexisting medical conditions can predict to some extent whether an individual will become severely ill or not, but the prediction is not very accurate. The early phase of infection results in direct tissue damage, followed by a late phase when the infected cells trigger an immune response, by recruitment of immune cells that release cytokines (reviewed in [1]). In severe patients, this may result in a “cytokine storm” and a systemic inflammatory response. Many individuals do not respond well enough to the vaccine, either because of old age or immune impairments. Thus, there is an ongoing search for anti-viral therapies and passive vaccines, as well as research into the basic mechanisms related to the virus and immunity towards it.

One useful path to investigate the immunity towards SARS-CoV-2 is adaptive immune receptor repertoire sequencing (AIRR-seq) [2, 3, 4], revealing noticeable changes in affected individuals in many arms of the immune system [5, 6]. Millions of B and T cell receptor (BCR and TCR, respectively) sequences from hundreds of individuals have been shared in public archives such as iReceptor [7] and OAS [8]. Thousands of individual antibody sequences validated as targeting and neutralizing SARS-CoV-2 have been published in datasets such as CoV-AbDab [9].

In the past few years, several studies have used AIRR-seq data to train machine learning (ML) algorithms to classify individuals who carry diseases [10], including celiac [11, 12], hepatitis C virus infection [13], cytomegalovirus [14], and others [15]. Finding the connection between AIRR-seq data and health states is a highly challenging task, because of the massive volume of AIRR-seq datasets that can include tens of millions of sequences that dilute the disease-specific biological signals. Another difficulty is our inability to determine to which antigen(s) each receptor can bind based solely on the receptor sequence. New methods to identify relevant repertoire features are continuously developed [10, 16, 17]. Besides the diagnostic and prognostic potential, such features can be critical in teaching us about the mechanisms behind the disease and the successful immune response towards it. Thus far, the vast majority of efforts to classify the health state or severity of COVID-19 have relied on TCR data [18, 19, 20, 21]. Recently, for example, a new approach to detect SARS-CoV-2 infection by TCR sequencing has been FDA approved for clinical use [20].

B cells undergo affinity maturation after pathogen encounter, to further adapt to the specific pathogen. Affinity maturation includes iterative cycles of somatic hypermutation (SHM) and affinity dependent selection. While selection depends on better binding, the SHM mechanism is independent of pathogen affinity. During SHM, different enzymatic pathways orchestrate together to introduce mutations specifically in the genomic regions encoding the antibody [22]. Extensive investigations have been devoted to understanding the SHM mechanism [23, 24, 25, 26], but to the best of our knowledge, no connection of a specific infection to a specific SHM pathway or pattern was made. The use of BCR sequencing is considered more difficult than TCR, because of SHM and higher diversity in the complementary determining region 3 (CDR3). It has been reported that BCR sequencing data cannot be used to classify individuals with COVID-19 [21]. Nevertheless, BCR data may be more informative than TCR in some cases, as BCRs undergo affinity maturation to adapt to each pathogen.

Here, using bulk and single cell BCR sequencing data, we successfully classify SARS-CoV-2 infected vs. naive individuals, as well as determine disease severity. Compared with the traditional sequence similarity clustering based approach, we obtain better classifications by considering SHM pattern changes in SARS-CoV-2 infected individuals. SHM specific patterns connected to decreased severity, as well as important amino acid (AA) composition in SARS-CoV2 antibodies, were identified.

## Methods

### Collection of samples

The repertoires composing the dataset were collected at three medical centers. IRB approval numbers: Rabin (Beilinson) Medical Center, 0256-20-RMC; Baruch Padeh Medical Center, 0037-20-POR; Shaare Zedek Medical Center, 0303-20-SZMC. 28 samples of controls were collected, as well as 39 mild patients with COVID-19 and 12 severely infected patients. Patients’ data can be found in Table S1. We do not have information about the SARS-CoV2 strains, but they are almost certain to be the original strain (before Alpha (B.1.1.7)). All samples were collected between April and early November 2020, and the earliest documented variant strains, as well as the earliest vaccines, arrived in Israel in late December 2020.

### Library preparation

#### Bulk

Ig repertoires were bulk sequenced according to the method described in [27]. All controls as well as 32 COVID-19 patients were sequenced for both heavy and light chains. These were used as the train/validation groups for the ML algorithms. For the rest of the patients, only heavy chains were sequenced, and served as the test group. 13 more controls for the test group were added from previously published datasets. Nine controls from dataset [28], and four from dataset [29].

#### Single cell

PBMCs from 13 individuals were prepared from fresh 5ml blood samples, and frozen according to the manufacturer’s instruction of the “Fresh Frozen Human Peripheral Blood Mononuclear Cells for Single Cell RNA Sequencing” protocol, document number CG00039 Rev D, 10X Genomics. Patients’ data can be found in Table S2. We do not have information about the SARS-CoV2 strains, as these tests were not routinely performed at that time (January-February 2021). Patients were not vaccinated. Libraries were prepared according to the manufacturer’s instruction of the “Chromium Next GEM Single Cell 5’ Reagent Kit v2 (Dual Index)” protocol, document number CG000331 Rev A, 10X Genomics. Libraries were pooled, mixed with 1% PhiX, and sequenced on an Illumina NovaSeq twice using an SP and an S1 kits.

### Data processing and statistics

FASTA files were generated using the PRESTO pipeline [30], and aligned to IMGT IGHV/D/J genes [31] using the VDJbase pipeline. Only sequences which started at the first 30 bases of the V gene were included. Isotype frequencies, V, D, J and combinations of V & J gene usage and CDR3 AAs 3-mers, as well as CDR3 AA lengths and V gene identities were calculated using a custom-designed R script (see data and code availability section). The same script also calculated the frequencies of BCR clusters (sharing the same V and J genes and junction AA length). Diversity was calculated using the alphaDiversity function from the Alakazam R package [32]. All P values were calculated using Wilcox test and adjusted using the Benjamini-Hochberg procedure [33].

### Generating an SHM model

A 5-mer SHM model was built using the function createTargetingModel from the shazam R package [23], once for silent mutations only and once for both silent and replacement mutations. To create these metrics for one representative from each clone, we used the collapseClones function from the same package. For each repertoire, substitutions, mutability, and targeting values were collapsed into a single table. Tables from all repertoires were collapsed into a single table. The tables enable both training ML algorithms and calculating mean mutability in specific sites (WR<u>C</u>/GTW and W<u>A</u>/<u>T</u>W hot-spots, the SYC/GRS cold-spot and all other sites). The table was also used to calculate single base mean mutability levels in all repertoires. The single base mutability was calculated as the average of all 5-mers with the same base in the middle.

### Training and estimation of ML algorithms

50 random splits to train and validation groups were made in order to estimate the F1 score, accuracy, sensitivity, and specificity of each model. Lasso and Elastic-Net Regularized Generalized Linear Models (GLMNET) using the caret R package [34] were trained on tables containing data from the repertoires. Feature selection was done using t-test calculations between frequencies in the different groups in the train subset only. Only features with P value below a certain threshold were selected. The algorithm was then trained on the selected data, and classifications were made for the validation groups. F1 score, accuracy, sensitivity, and specificity were calculated for each random split.

### COVID-19 classification using AA frequencies at all V gene positions

Frequencies of each AA along 103 positions (according to the IMGT numbering) in each V gene family were calculated for all repertoires. The train/validation samples were used to train the same algorithm as explained above, and to estimate the F1 score, accuracy, sensitivity, and specificity of the algorithm. The validation group was used to estimate the parameters of the algorithm on unseen data. Coefficients of the algorithm were extracted and enabled to calculate scores for single antibodies. If a certain AA was present in the sequence, it received a frequency of 1. Otherwise, it received a frequency of 0. This equation was used to calculate scores for all antibodies in all repertoires, as well as scores for known COVID-19 antibodies from the CoV-AbDab database.

### Single cell data analysis

Single cell data was analyzed using cell-ranger 6.0.1 with output of both VDJ recombination and gene expression data. Cell-ranger output was then manipulated using the Seurat R package [35]. Cells with more than 5% mitochondrial gene expression were removed. Data was normalized, and PCA and UMAP on the top 10 PCAs were done using standard Seurat functions. Cell identity was determined using the SingleR R package against a sorted dataset from the celldex R package [36]. Barcodes of VDJ data and gene expression data were matched using R.

## Results

### BCR gene usage cannot classify SARS-CoV2 infection

To assess changes in BCR repertoires of COVID-19 patients, we collected 79 blood samples and sequenced their BCR repertoires. Samples were split to three groups: uninfected individuals, mildly and severely COVID-19 infected patients. For each group we characterized several whole repertoire features, such as CDR3 AA length distribution, V gene mutation distribution, clonal diversity, V, D, J and combination of V and J gene usage. We also calculated frequencies of BCR clusters (same V and J gene as well as same CDR3 AA length). These measurements are shown in Fig. 1 and in Fig. S1 for heavy chains, and for kappa and lambda light chains in Figs. S2 and S3. As expected, the diversity of BCR clones is significantly lower in COVID-19 patients compared with controls (Fig.1C). No significant difference was observed in CDR3 AA length (Fig.1A), and only slight increase was seen in V gene mutation distribution (Fig.1B). For many V genes we observed significantly reduced usage in COVID-19 patients (Fig.1D). Three exceptions are IGHV4-34, IGHV4-39 and IGHV4-59 that demonstrate increased usage upon infection, which is further increased in severe patients compared with mild ones. These results support previously published COVID-19 data [37, 38], and suggest that antibodies against SARS-CoV2 mainly comprise those genes. To further validate these conclusions, we tried to build ML classifiers based on V, V & J gene usage, or V & J gene usage and 85% similarity in the CDR3 AAs. However, these models yielded less than 70% accuracy, suggesting low impact of V or V & J gene usage on the response to SARS-CoV2 infection.

**Figure 1:**
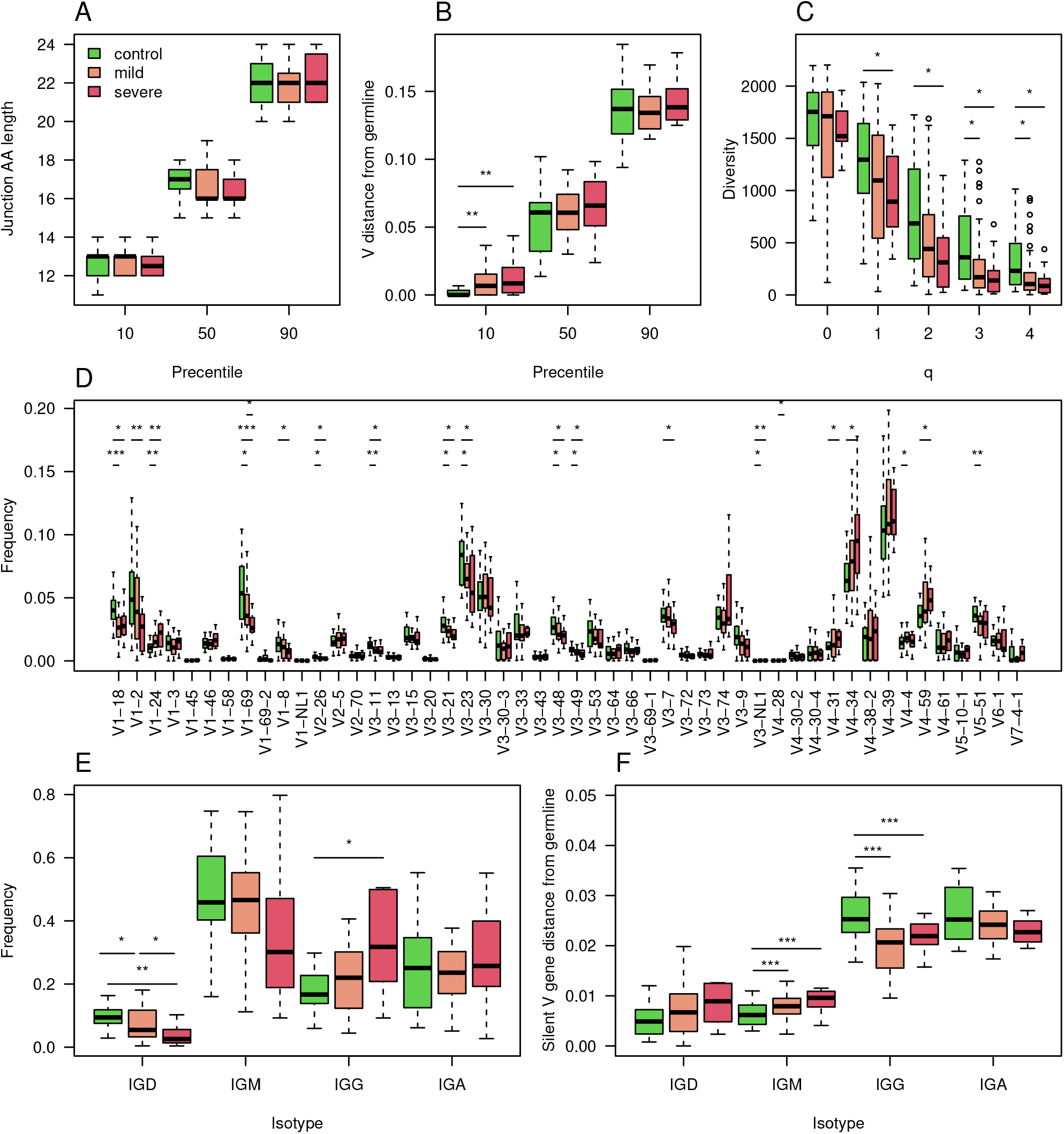
Characterization of the COVID-19 heavy chain BCR cohort. A. 10,50 and 90 percentiles of AA CDR3 length in individuals with corona at indicated severity and controls. B. 10,50 and 90 percentiles of V gene distances from germline in COVID-19 infected individuals at indicated severity and controls.C. Boxplot showing calculated Hill diversity indexes upon different q values between individuals infected by COVID-19 at indicated severity and controls. D. Boxplots showing V gene usage in individuals infected by COVID-19 at indicated severity and controls, shown top 50’s mean frequencies. E. Boxplots showing the isotype frequencies in individuals infected by COVID-19 at indicated severity and controls. F. Boxplots showing silent mutations’ frequencies along the V gene in different isotypes of individuals infected by COVID-19 at indicated severity and controls. In the whole figure, * marks P value less than 0.05. ** marks P value less than 0.01 and *** marks P value less than 0.001.

We explored further whole repertoire features, and compared isotype frequencies between the different groups. While we observed a reduction in the frequencies of IGD and IGM upon SARS-CoV2 infection, the levels of IGG increased (Fig.1E), and those of IGA remained unchanged. We also measured silent mutability frequencies for each isotype (Fig. 1F). These measurements avoid changes which are caused by antibodies selective pressure. In contrast to the IGG and IGA class switched isotypes, in which mutability upon infection is reduced, in IGD and IGM mutability is increased. In severe patients, the IGD and IGM mutability was even higher (Fig.1F).

### BCR V gene AA composition successfully classifies SARS-CoV2 infection and may reveal important features of antibodies against the virus

We continued exploring classification approaches to stratify COVID-19 patients and uninfected individuals. To this end, we explored AA frequencies along the V gene, aggregated by V gene family. We generated a table with 10,300 columns, counting AA frequencies along 103 V gene positions (aligned according to IMGT numbering), for the 5 most highly used V gene families (IGHV1-5). Using this approach we obtained a high F1 score of more than 0.85, and similar levels of accuracy, sensitivity,and specificity (Fig.2A). The test set resulted in an F1 score of above 0.85 (Fig. 2B). We then extracted the coefficient used by the algorithm, corresponding to the contribution of each AA frequency to the classification of the disease (Fig. 2D).

**Figure 2:**
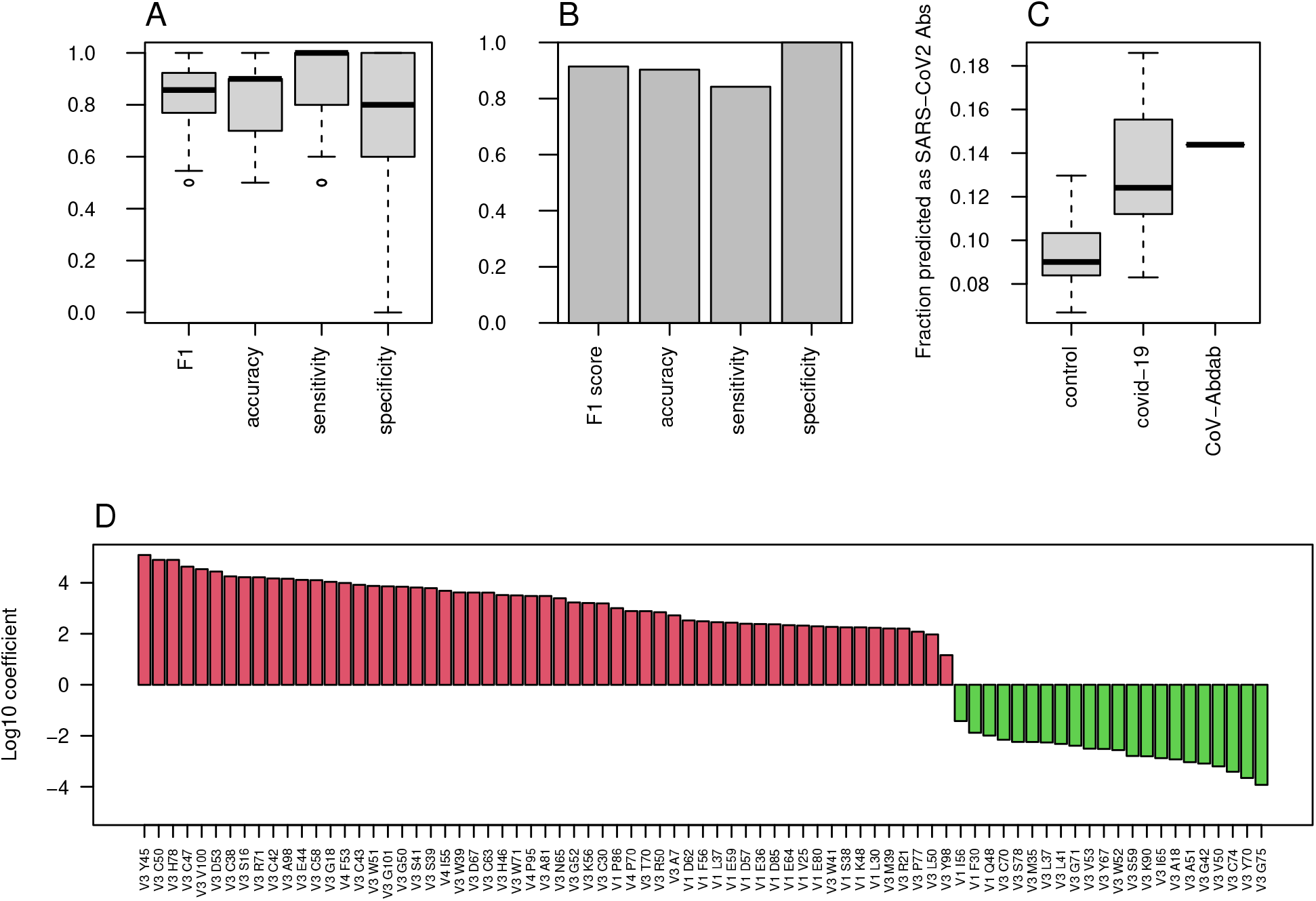
COVID-19 classification using AA frequencies at all V gene positions. A. Boxplots showing the F1 score, accuracy, sensitivity, and specificity for COVID-19 classification by AA frequency at each position in each V family. Shown are values calculated for 50 random splits to train and validation groups. B. Bar plots showing the indicated scores on the external test group. C. COVID-19 single antibody scores were calculated using the coefficients of the algorithm described in panel A. Boxplos showing the fraction of antibody sequences with scores above 0 in control and COVID-19 infected repertoires, as well as in CoV-AbDab COVID-19 antibodies, are shown. D. Log10 coefficients of the algorithm described in A and B.

To further validate that these changes are unique to COVID-19 patients, we downloaded a dataset of more than 450 repertoires from cAb-rep data collection [39]. These data include repertoire sequencing results from a wide variety of clinical conditions such as Hepatitis B virus infection, vaccinations against Hepatitis B virus and influenza, and several autoimmune diseases. Applying our algorithm to these data to classify COVID-19 infection resulted in a false positive rate of only 6%, indicating that our classification is specific to COVID-19 infection.

These results were obtained for the repertoire level, and we sought to test their applicability to the single BCR sequence level. For this, we transferred the features selected for the repertoire level model, i.e., AA frequencies along the V gene families, to calculate a score for single BCR sequences. We calculated such scores for a list of more than 5,000 known antibodies against SARS-CoV2 from the CoV-AbDab database [40]. The scores of the known antibodies were higher than those came from whole repertoires of control patients as well as most of the COVID-19 infected repertoires (Fig. 2C), suggesting that these coefficients are meaningful not only for the repertoire level, but also for single BCR sequences. Our attempts to classify the severity of COVID-19 using this method were not successful, so for this purpose, we explored other sets of features. The coefficients of the algorithm can be seen in Fig. 2D.

### Mutation bias in class-switched B cells of COVID-19 patients

As reduced levels of overall BCR mutability were seen upon SARS-CoV2 infection only in the class switched isotypes (Fig 1F), we quantified single base mutability patterns in these isotypes. As seen in figure 3A, the mean relative mutability is reduced in COVID-19 patients at Cytosine and Guanine (C and G), but increases in Adenine and Thymine (A and T). The same results were obtained when considering silent mutations only (Fig. 3B). Five main pathways are responsible for introducing mutations during SHM [41]. Three introduce mutations in C and G, and the other two involve the low fidelity DNA polymerase pol*η*, which mutates A and T. The significant differences in mutability observed in COVID-19 patients suggest altered activity of those arms. To further investigate SHM in SARS-CoV2 infection, we applied a commonly used 5-mers SHM mutability model [23]. In general, two highly mutated hot-spot motifs are commonly observed in SHM. One is WR<u>C</u>/<u>G</u>YW (where W = {A, T}, Y = {C, T} R = {G, A}, and the mutated position is underlined), and the other is W<u>A</u>/<u>T</u>W. In addition, SY<u>C</u>/<u>G</u>RS (where S = {C, G}), is considered as a cold-spot sequence motif. We first built a 5-mer mutability model based on both silent and replacement mutations. Such a model combines the effects of SHM and antigen-driven selection. We divided the 5-mers to those occurring in the two hot-spots, in the cold-spot, and in all other neutral sites, and show their levels for IGD/IGM and for IGA/IGG (Fig. 3C and E). The most significant changes between the different groups are a decrease in the WR<u>C</u>/<u>G</u>YW site and an increase in SY<u>C</u>/<u>G</u>RS in IGA/IGG of COVID-19 patients. This increase is not seen in severely infected patients.

**Figure 3:**
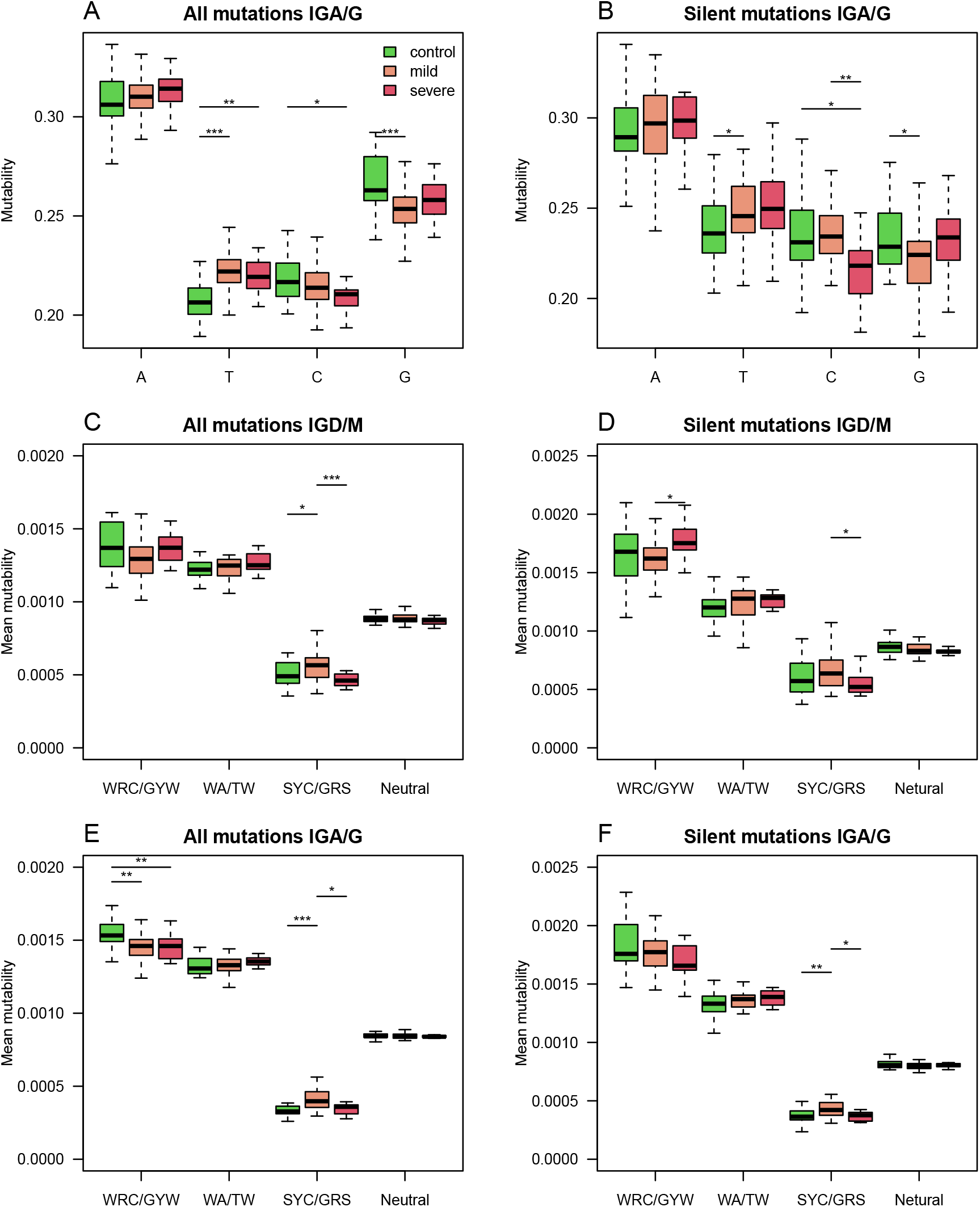
Silent and replacement mutability in SHM single base mutability, 5-mers hot-spots and cold-spots. A single base mutability model was built based on IGA/G isotypes of COVID-19 patients and controls. Shown are boxplots representing the normalized sum of single base mutability. The same plot as in A but for silent mutations only. C-D. An 5-mer SHM model based on both silent and replacement mutations in C, or silent only mutations in D, was built using the IGD and IGM isotypes of COVID-19 patients at different severity levels and controls. Shown mutability of the two known SHM hot-spots, SHM cold-spots, and the rest of the sites. E-F. An 5-mer SHM model based on both silent and replacement mutations in E, or silent only mutations in F, was built using the IGA and IGG isotypes of COVID-19 patients at different severity levels and controls. Shown mutability of the two known SHM hot-spots, SHM cold-spots, and the rest of the sites. In the whole figure, * marks P value less than 0.05. ** marks P value less than 0.01 and *** marks P value less than 0.001.

To understand whether these patterns stem from SHM or from antigen-driven selection, we built another model, taking only silent mutations into consideration. Fig. 3D and F shows the resulting mutability scores for the same sequence motifs. The observed pattern resembles the one observed in Fig. 3C and E, suggesting that the alteration between the groups results from altered SHM characteristics. To avoid the effect of clonal expansion on mutability calculations, we repeated all calculations, taking into account only one representative from each clone. Similar results were obtained using this approach (Fig. S4). Moreover, using SHM matrices based only on a specific V family resulted in a much lower signal (Fig. S5F). Importantly, the mentioned SHM patterns reflect the relative likelihood for each mutation pattern and do not indicate the overall mutability level.

### Silent SHM patterns classify SARS-CoV2 infection and severity

To estimate the level of connection between changes in SHM patterns and SARS-CoV2 infection, we tried again to build a classifier of samples’ origin. We built two models, one using all mutations (Fig. 4A, S5, S6A and S8), and one using silent mutations only (Fig. 4B, S6B). Taking all mutations into account, we obtained an F1 score of over 0.85, as well as accuracy, sensitivity, and specificity values. Taking only silent mutations into account, we obtained a slightly lower result of ∼ 0.8 F1 score and accuracy. These results strengthen our hypothesis that the differences between the repertoires emerge mainly from SHM itself and not from antigen-driven selection. Using only light chain sequences for the mutability model reaches much lower results, as expected (Fig. S7A and B). A model based on the combination of light and heavy chains does not obtain better results than using the heavy chain only (Fig. S8).

**Figure 4:**
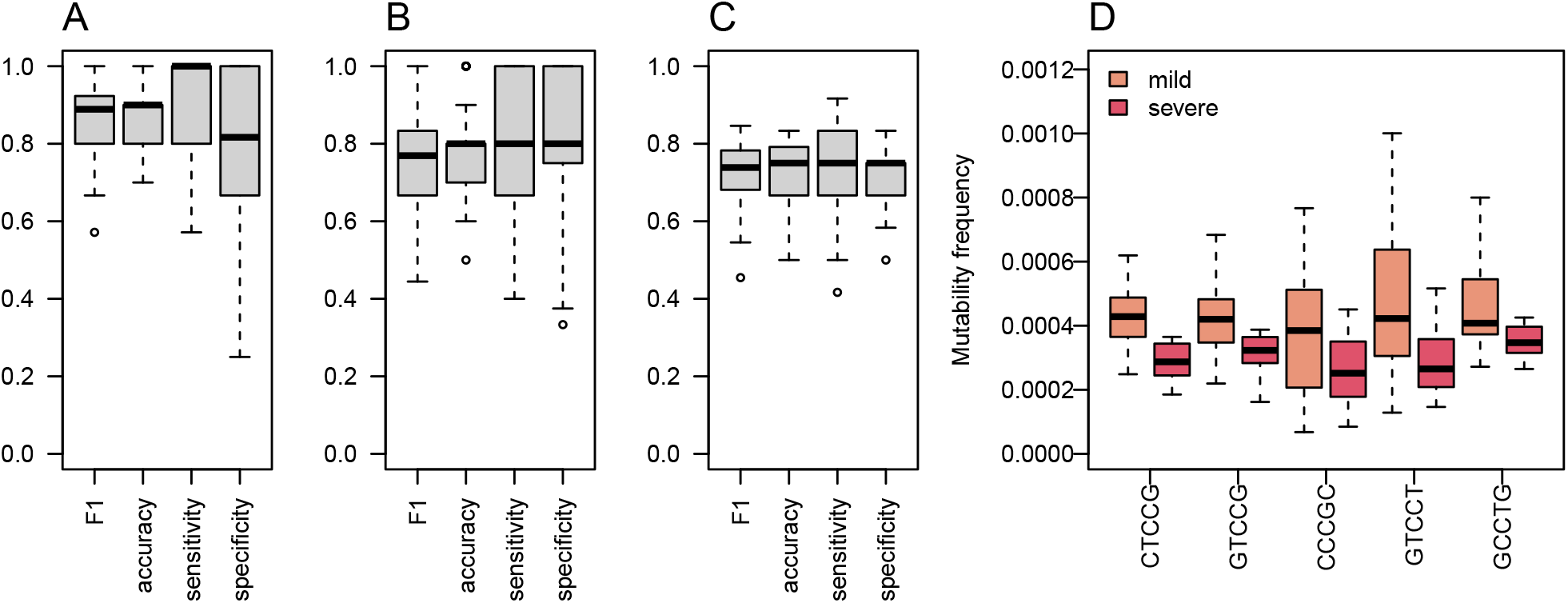
SHM Heavy chain enables classification of both SARS-CoV2 infection and COVID-19 severity. A. An ML algorithm was trained on the substitutions matrix of the 5-mer SHM model, which was created for the IGA/G isotypes. Boxplots representing F1 score, accuracy, specificity, and sensitivity of 50 random splits to train and test groups are shown. B. The same algorithm as in A was trained on silent mutations only. Shown are Boxplots representing the F1 score, accuracy, specificity, and sensitivity of 50 random splits to train and test groups. C. Boxplots showing F1 score, accuracy, specificity, and sensitivity of 20 leave-one-out cross validation of severity classification. Each leave-one-out was on 12 severe COVID-19 patients and 12 randomly selected mild COVID-19 patients. The ML algorithm was trained on the mutability matrix of the SHM cold-spots in these groups. D. Frequency of mutability in mild and severe individuals with COVID-19. Boxplots of frequencies of repeating coefficients of the algorithm explained in C are shown.

Next, we tried to classify COVID-19 severity using SHM patterns. Since the mutability in the cold-spot motif changes the most between severe and mild patients, we built a model using mutability scores of this cold-spot only. We obtained an F1 score and accuracy of about 0.75 in severity classifications (Fig. 4C).

All patterns with non-zero coefficients have much higher mutability frequencies in mild patients compared with severe patients ((Fig. 4D). Again, to avoid the effect of clonal expansion and selective pressure on the inferred mutability model, we repeated the mutability model inference taking into account only one representative from each clone. As shown in Fig. S5, the results were comparable to those obtained using all sequences.

### Known SARS-CoV2 antibodies are enriched in plasmablasts from COVID-19 patients

We thought to find in our sequencing data, antibodies that may be related to the known COVID-19 antibodies. As mentioned above, during the COVID-19 pandemic a new database summarizing all known SARS-CoV2 antibodies was published, containing more than 5,000 antibody AA sequences of both heavy and light chains. For each of our repertoires, we calculated and summarized the frequencies of sequences that are similar to known antibodies. We defined similar antibodies by 85% identity in the CDR3 AAs, and the same V and J genes. As expected, the frequencies of similar to known antibodies in COVID-19 patients were higher than those in control individuals (Fig. 5A. Histograms summarizing the sizes and numbers of samples having at least one representation in the clones can be found in Fig. S9A and B). Using the sum of frequencies of similar to known COVID-19 clones, we reached an accuracy of above 70% in repertoire classification and an AUC of 0.81 (Fig. 5B). Even lower results were obtained when training the algorithm to count the frequencies of shared clones between samples (Fig. S10). Although significant, this result is lower than that achieved by considering mutations along the V gene.

**Figure 5:**
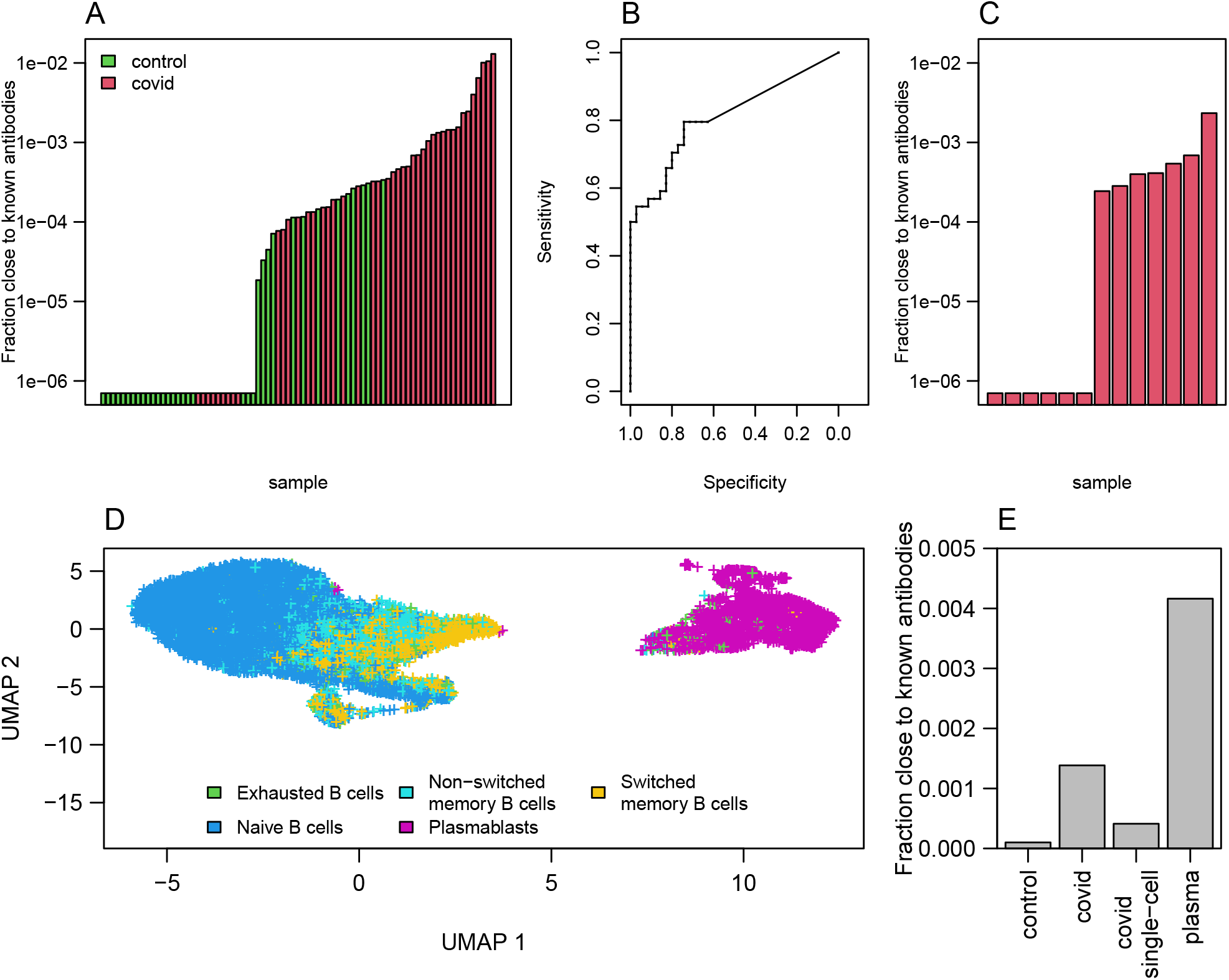
Clones of antibodies in our sequencing close to known COVID-19 antibodies from CoV-AbDab database. A. Sum of frequencies of clones (same V and J genes and 85% similarity in AA of CDR3) close to known COVID-19 antibodies (from CoV-AbDab data base) in COVID-19 patients and controls. B. ROC curve summarizing the results shown in A. C. Sum frequencies of clones close to COVID-19 antibodies in 13 single cell COVID-19 patients data. D. UMAP on gene expressions of B cells isolated from 13 patients showing differences between naive, memory and plasmablast cells. Cell type identification was done using SinglR. E. Sum of frequencies of antibodies close to known COVID-19 antibodies in bulk sequencing of COVID-19 patients and control as well as in sequences from single cell sequences of COVID-19 patients and in cells identified as plasmablast cells.

To further explore the similarity to known antibodies, we performed 10X Genomics single cell sequencing including V(D)J and gene expression, on blood samples from additional 13 mild COVID-19 patients. Using single cell sequencing data enables matching of heavy and light chains, which cannot be done with bulk sequencing. Moreover, single cell sequencing provides the ability to identify cell type using gene expression signatures. We found similar to known antibodies in 7 out of the 13 repertoires. The frequencies were overall lower compared with those seen in the bulk RNA sequencing cohort (Fig. 5C). This could be due to the differences in sequencing methods, or because in the single cell cohort the patients were diagnosed on average more recently than the bulk cohort and thus may have had lower levels of SARS-CoV2 specific antibodies.

We then applied the SingleR R package to classify cell types by single cell expression profiles. Two-dimensional UMAP reduced plots are shown in Fig. 5D, demonstrating a distinct cluster of plasmablasts. We summarized the frequency of known SARS-CoV2 clusters in bulk sequenced COVID-19 patients, bulk controls, single cell unsorted data, and single cell plasmablasts only. As shown in Fig. 5E, COVID-19 patients show enriched levels of similarity to known SARS-CoV2 antibody compared with controls. Single cells show higher levels than controls but lower than bulk, as discussed above. Among plasmablasts of COVID-19 patients, we see the highest frequency of known antibody clusters, indicating a stereotypical response to SARS-CoV2. Lastly, to validate our observation that WR*C*/<u>G</u>YW hot-spots mutability scores decrease upon COVID-19 infection, and SY<u>C</u>/<u>G</u>RS cold-spots increase (Fig. 3), we split the single cell data into plasmablasts vs. all other B cell types. We built a mutability SHM matrix for each of these subsets, and indeed found a reduction in the mutability scores of WR<u>C</u>/<u>G</u>YW hot-spots in plasmablasts (0.00168) compared with the other B cell types (0.00178), and an increase in the mutability scores of the SY<u>C</u>/<u>G</u>RS cold-spots (0.0003 and 0.0002, respectively).

## Discussion

The COVID-19 pandemic, caused by evolving variants of SARS-CoV2, has infected a large proportion of the population worldwide. Antibodies play a critical role in eliminating the virus from the body. Serological tests are routinely used to estimate immunity of individuals against SARS-CoV2, convalescent plasma donations were used to treat severely ill COVID-19 patients, and many monoclonal antibodies were developed as candidate passive vaccinations.

Although the pandemic has caused a huge health and economic burden, it brought several important advantages for biomedical research. With so many researchers and funding opportunities focusing on a single topic, the pandemic facilitated both broad and profound analyses of the virus and the immune responses towards it. During the past two and a half years, thousands of COVID-19 binding/neutralizing antibodies have been published and deposited in public datasets[42, 43]. This huge amount of data facilitates finding BCR sequences that are similar to known antibody sequences, and searching for common features. Such features may be used in the clinic for diagnosis of the disease, but in the case of COVID-19 there are easier, faster and cheaper ways to do that. Much more importantly, it can teach us about the development of the immune response towards the virus.

Here, in contrast to previous reports[21], we were able to stratify COVID-19 patients and healthy individuals based on shared clusters of BCR sequences. The moderate classification results of such approach led us to explore different sets of features that turned out to be more informative. AA frequencies at all V gene positions served as a basis for an ML model that produced a high F1 score (∼ 85%) in classifying COVID-19 infection.

The patterns of AA alterations in BCRs arise during the process of affinity maturation, that includes two iterative processes, namely SHM and affinity-dependent selection. These patterns can stem from the antibodies against SARS-COV2 or from overall altered SHM mechanism in COVID-19 patients.

An important question that may arise when inspecting the presented approach is whether it is specific to COVID-19, or perhaps it simply detects general signals related to an adaptive immune response towards a new pathogen. We believe that the presented approach is specific to COVID-19 because: 1. The signal does not disappear when choosing a single representative per clone, which eliminates the effect of general clonal expansion. 2. The signal is based on an SHM pattern, which is subject to an antigen-specific affinity maturation. 3. Our lab has a lot of experience in ML-based classification of different clinical conditions[44, 17, 28], and for each condition the features identified by the algorithm as the most essential for classification were different. SHM patterns have never been previously identified as a feature, as far as we know (but see our recent publication [45]). To test this, we applied our algorithm to data from ∼450 samples, including infection with Hepatitis B virus, vaccinations against Hepatitis B virus and influenza, and several autoimmune diseases. 94% of these repertoires were classified as healthy, indicating that our algorithm does not classify any neo-response as COVID-19.

Extensive research has been devoted to study SHM mechanisms affecting other regions in the antibody besides the CDR3[46, 23]. Yet, this knowledge has not been used for disease classifications, nor for improving antibody engineering. We sought to follow the SHM machinery during SARS-CoV2 infection, starting with the whole repertoire level. It is well established that antibodies binding SARS-CoV2 are very close to the germline[47, 5, 48, 49]. Surprisingly, even at the repertoire level, we detected a decrease in mutability of IGG BCRs. To explore whether the AA frequency-based signal results from alterations in SHM or affinity dependent selection, we followed the mutability rates of silent mutations only. These mutations are not subjected to affinity dependent selection pressure, thus reflecting changes in the machinery of SHM. We found that most SHM changes upon SARS-CoV2 infection were observed even when counting only silent mutations, which are not subject to affinity selection, suggesting dramatic changes in the SHM machinery upon SARS-CoV2 infection. To further pinpoint the effects on the SHM machinery, we repeated the calculations taking only one representative from each clone into account, thereby abolishing the effect of clonal expansion (Fig. S5). This step slightly reduced the F1 score, in a non-significant way. The fact that eliminating the effect of clonal expansion on our findings did not abolish the differences suggests that there are true changes in the SHM machinery. Moreover, the moderate performance reduction when taking only one representative per clone, hints that the SHM changes during SARS-CoV2 infection may be further enhanced by clonal expansion, potentially aiding the battle with the virus.

Many pathways are involved in the introduction of mutations to BCR sequences. In particular, two common SHM hot-spots, WR<u>C</u>/<u>G</u>YW and W<u>A</u>/<u>T</u>W, are affected by two different pathways. While mutations in WR<u>C</u>/<u>G</u>YW motifs are mediated by the activation induced deaminase, mutability at W<u>A</u>/<u>T</u>W motifs also involve the low fidelity DNA polymerase pol*η*.

In the class switched IGA and IGG isotypes, we observed decreased mutability levels with increasing severity of COVID-19 at WR<u>C</u>/<u>G</u>YW motifs, and increased mutability at W<u>A</u>/<u>T</u>W sites. Again, these changes were observed even when counting silent mutations only, further supporting an impact of the virus on the SHM introduction mechanism. The reduced mutability in WR<u>C</u>/<u>G</u>YW motifs and the mildly increased mutability in W<u>A</u>/<u>T</u>W motifs may hint that AID levels could be decreased upon COVID-19 infection. This possibility will need to be validated in future studies. Another future direction is to test for possible SHM positional effects. The presence of such an effect was lately suggested [50], and it will be very interesting to inspect whether this is relevant to our results.

Another specific SHM target is the cold-spot SY<u>C</u>/<u>G</u>RS. Surprisingly, we found an increase in mutability rates of this cold-spot in COVID-19 repertoires. Moreover, this increase was not observed in severely infected patients, suggesting that this mechanism may be critical for production of efficient antibodies and thereby for prevention of severe illness.

Building on our success in classifying patients from healthy individuals, we sought to develop an ML-based algorithm to classify disease severity. This could have important clinical outcomes, since medications and passive vaccines now exist that can prevent deterioration if diagnosed individuals are treated rapidly. However, these treatments have side effects and are not given to the wide population. Prediction of disease severity by the known risk factors is highly inaccurate, and there are currently no other means to classify severity. Using mutability patterns from silent mutations only, we estimate our ability to classify COVID-19 severity at approximately 75%(Fig. 4C). The known risk factors to develop severe COVID-19 are mostly preexisting conditions such as older age, hypertension, obesity, diabetes. Here, we suggest another risk biomarker that involves basic features of the adaptive immune system. Many more steps are needed to enable prediction of COVID-19 infection and severity based on BCR sequencing data. We provide here a first step towards it.

AA frequency patterns along the V genes at the whole repertoire level is a sufficient feature for relatively good classification of COVID-19. Looking at the identity of AA along the V gene of a single BCR sequence may reveal its affinity towards the virus. To explore the connection between the new BCR repertoire data generated here and known SARS-CoV2 antibody sequences we took a two way approach. Building on the hypothesis that the whole repertoire level signal responsible for the classification stems from individual SARS-CoV2-specific antibodies generated during the infection, we derived a single sequence score based on the repertoire classification signal. Although sequences with high scores are scarce in both healthy and COVID-19 repertoires, their prevalence in the CoV-abDab data is significantly higher (Fig. 2C). As such, the features (detailed in Fig. 2D) may be used for more rational antibody design towards the virus. In addition, we explored the presence of similar sequences to the validated CoV-abDab antibodies in both bulk and in single cell sequenced repertoires. We found a higher fraction of sequences with high similarity to known antibodies in COVID-19 patients compared with controls. This can also be used for successful classification of the repertoires. Notably, a group of COVID-19 patients had no similar antibodies to those in the list, suggesting that despite the massive efforts so far, the list is incomplete. On the other hand, in some control samples we found few sequences similar to known antibodies. These antibodies may provide a basis for protection from COVID-19 symptoms or complications to individuals who carry them.

## Declarations

### Ethics approval and consent to participate

The repertoires composing the dataset were collected at three medical centers. IRB approval numbers: Rabin (Beilinson) Medical Center, 0256-20-RMC; Baruch Padeh Medical Center, 0037-20-POR; Shaare Zedek Medical Center, 0303-20-SZMC. All participants received an explanation about the study from a medical doctor, and signed an informed consent form.

### Consent for publication

Not applicable.

### Availability of data and code

Our sequencing data will be available on NCBI upon publication, under BioProject PR-JNA839749. All code will be available on github.

### Authors’ contributions

GY, MGT, and TT conceived the research; GY supervised the work; MS performed the computational analyses; ZT prepared and sequenced the BCR libraries; PP coordinated between all the parties and transferred the samples from the hospitals to the lab at Bar Ilan University; SS, MM, HK, YH, AP, EBC, BB collected the samples from COVID-19 patients; ALB, VY collected the samples from healthy volunteers; MS, PP, GY wrote the manuscript; all authors edited the manuscript and approved it for publication.

## Acknowledgements

Not applicable.

