## Supplementary material for "Altered somatic hypermutation patterns in COVID-19 patients classifies disease severity"

Table S1: **Summary of volunteers' data (bulk sequencing).** All samples were collected in 2020.

| Hospital | Age | Sex | Condition | Diagnosis date | Sample date | Sample # |
|---|---|---|---|---|---|---|
| Rabin | NA | NA | mild | NA | 23.7 | HSCov1 |
| Rabin | 23 | F | mild | 30.7 | 2.8 | HSCov2 |
| Rabin | 58 | M | mild | 1.8 | 5.8 | HSCov4 |
| Rabin | 45 | M | mild | 21.7 | 9.8 | HSCov5 |
| Rabin | 55 | M | mild | 13.8 | 18.8 | HSCov6 |
| Rabin | NA | M | severe | 14.8 | 18.8 | HSCov7 |
| Shaare Zedek | 39 | M | mild | 22.4 | 23.4 | HSCov13 |
| Shaare Zedek | 31 | F | mild | 25.4 | 27.4 | HSCov14 |
| Shaare Zedek | 44 | F | mild | 28.4 | 30.4 | HSCov15 |
| Shaare Zedek | 43 | F | mild | 1.5 | 3.5 | HSCov16 |
| Shaare Zedek | 30 | F | mild | 15.5 | 17.5 | HSCov18 |
| Shaare Zedek | 59 | F | mild | 27.5 | 3.6 | HSCov19 |
| Rabin | 58 | M | mild | 26.8 | 26.8 | HSCov20 |
| Rabin | 68 | M | mild | 1.9 | 1.9 | HSCov21 |
| Rabin | 75 | M | severe | 30.8 | 1.9 | HSCov22 |
| Rabin | 61 | F | mild | 20.9 | 23.9 | HSCov24 |
| Rabin | 78 | M | mild | 17.9 | 24.9 | HSCov25 |
| Rabin | 61 | M | mild | 20.9 | 24.9 | HSCov26 |
| Rabin | 69 | F | mild | 23.9 | 30.9 | HSCov27 |
| Rabin | 49 | M | severe | 20.9 | 30.9 | HSCov28 |
| Rabin | 52 | M | severe | 27.9 | 1.10 | HSCov29 |
| Rabin | 57 | M | mild | 29.9 | 6.10 | HSCov30 |
| Rabin | 62 | F | severe | 4.10 | 6.10 | HSCov31 |
| Rabin | 86 | M | severe | 2.10 | 6.10 | HSCov32 |
| Rabin | 65 | M | mild | 29.9 | 7.10 | HSCov33 |
| Rabin | 69 | M | severe | 7.10 | 7.10 | HSCov34 |
| Rabin | 42 | F | mild | 14.10 | 21.10 | HSCov38 |
| Rabin | 18 | M | mild | 1.10 | 21.10 | HSCov39 |
| Rabin | 19 | F | mild | 3.10 | 21.10 | HSCov40 |
| poria | 53 | F | severe | 31.08 | 6.10 | HSCov43 |
| poria | 55 | F | mild | 30.08 | 4.10 | HSCov45 |
| poria | 49 | F | mild | 31.3 | 4.10 | HSCov49 |
| poria | 28 | M | mild | 19.9 | 4.10 | HSCov50 |
| poria | 47 | F | mild | 23.8 | 4.10 | HSCov51 |
| poria | 43 | M | severe | 4.10 | 6.10 | HSCov53 |
| poria | 48 | F | mild | 30.4 | 4.10 | HSCov56 |
| poria | 61 | F | mild | 28.3 | 6.10 | HSCov57 |
| poria | 26 | F | mild | 14.8 | 4.10 | HSCov58 |
| poria | 37 | M | mild | 30.7 | 4.10 | HSCov59 |
| poria | 51 | M | mild | 28.9 | 6.10 | HSCov60 |

Continued on next page

**Table S1 – continued from previous page**

| Hospital | Age | Sex | Condition | Diagnosis date | Sample date | Sample # |
|---|---|---|---|---|---|---|
| poria | 61 | M | mild | 25.9 | 6.10 | HSCov61 |
| poria | 34 | M | severe | 22.9 | 6.10 | HSCov62 |
| poria | 61 | M | severe | 5.10 | 6.10 | HSCov63 |
| poria | 58 | F | mild | 23.9 | 8.10 | HSCov64 |
| poria | 36 | M | mild | 24.9 | 8.10 | HSCov65 |
| poria | 27 | M | mild | 16.9 | 8.10 | HSCov66 |
| poria | 28 | F | mild | 2.9 | 8.10 | HSCov67 |
| poria | 55 | F | mild | 27.9 | 6.10 | HSCov68 |
| poria | 85 | M | mild | 21.9 | 6.10 | HSCov69 |
| poria | 34 | M | mild | 24.9 | 8.10 | HSCov70 |
| poria | 72 | M | severe | 2.11 | 3.11 | HSCov71 |

Table S2: **Summary of volunteers' data (single cell sequencing).** All samples were collected at Rabin Hospital, with a mild condition.

| Age | Sex | Diagnosis date | Sample date | Sample # |
|---|---|---|---|---|
| 31 | F | 27.12.20 | 30.12.20 | HSCov73 |
| 41 | F | 28.12.20 | 30.12.20 | HSCov75 |
| 39 | F | 29.12.20 | 24.1.21 | HSCov77 |
| 44 | M | 7.1.21 | 24.1.21 | HSCov78 |
| 26 | M | 23.1.21 | 24.1.21 | HSCov79 |
| 22 | F | 14.10.21 | 24.1.21 | HSCov81 |
| 45 | M | 10.10.21 | 31.1.21 | HSCov83 |
| 39 | F | 15.1.21 | 31.1.21 | HSCov85 |
| 58 | F | 14.1.21 | 31.1.21 | HSCov86 |
| 31 | F | 1.2.21 | 14.2.21 | HSCov87 |
| 43 | F | 3.2.21 | 14.2.21 | HSCov88 |
| 28 | F | 1.2.21 | 14.2.21 | HSCov89 |
| 49 | M | 14.1.21 | 14.2.21 | HSCov90 |

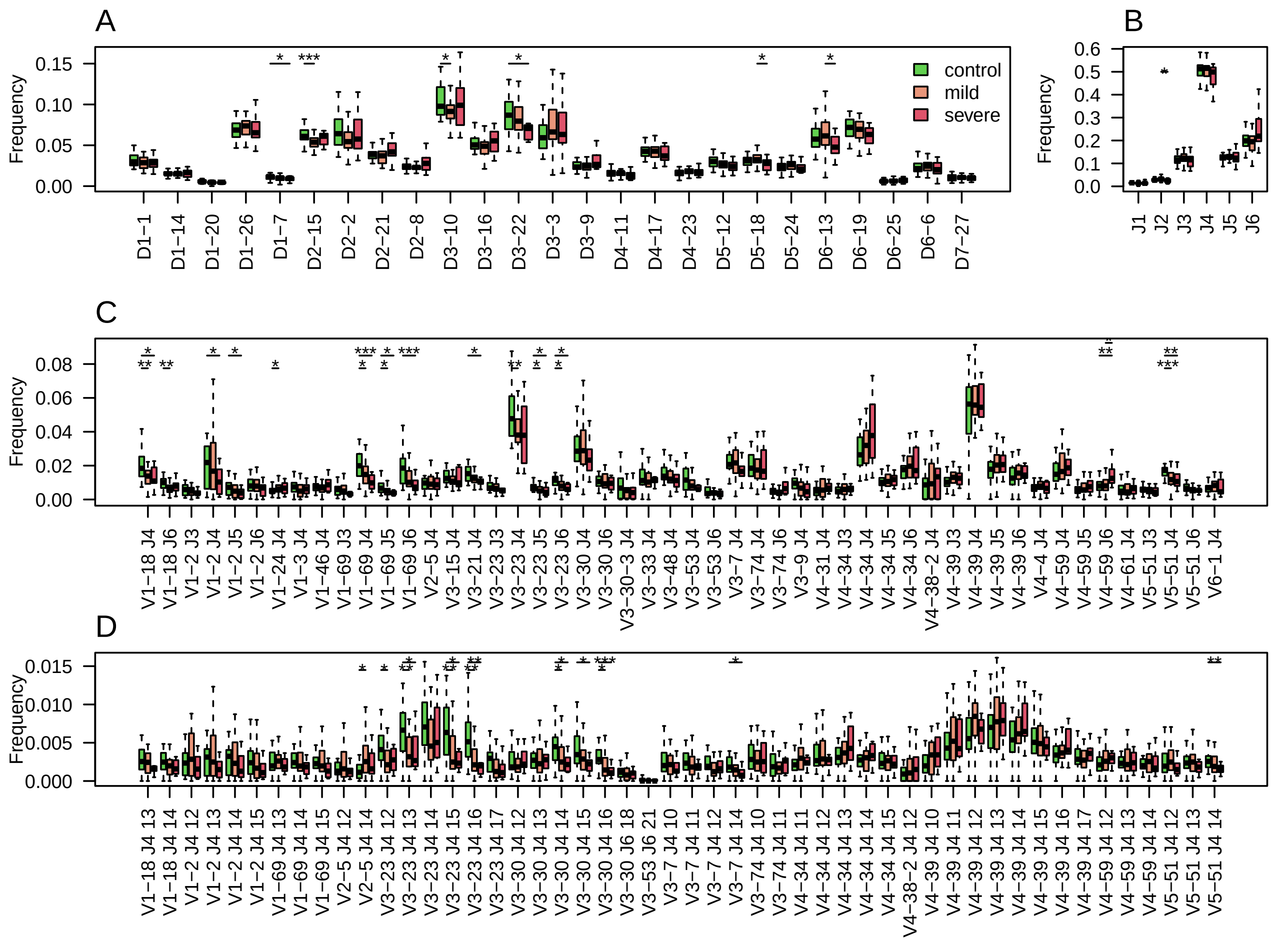


Figure S1: **Characteristics of the heavy chain sequencing data**
A. D gene usage comparison between individuals with COVID-19 at indicated severity levels and healthy controls. B. J gene usage comparison. C. Combinations of V & J gene usage comparison. Shown are the top 50 highest frequencies. D. Clusters comparison between individuals with COVID-19 at indicated severity levels and healthy controls. Shown are the top 50 highest frequencies. Throughout the figure, * marks a P value lower than 0.05, ** marks a P value lower than 0.01, and *** marks a P value lower than 0.001.

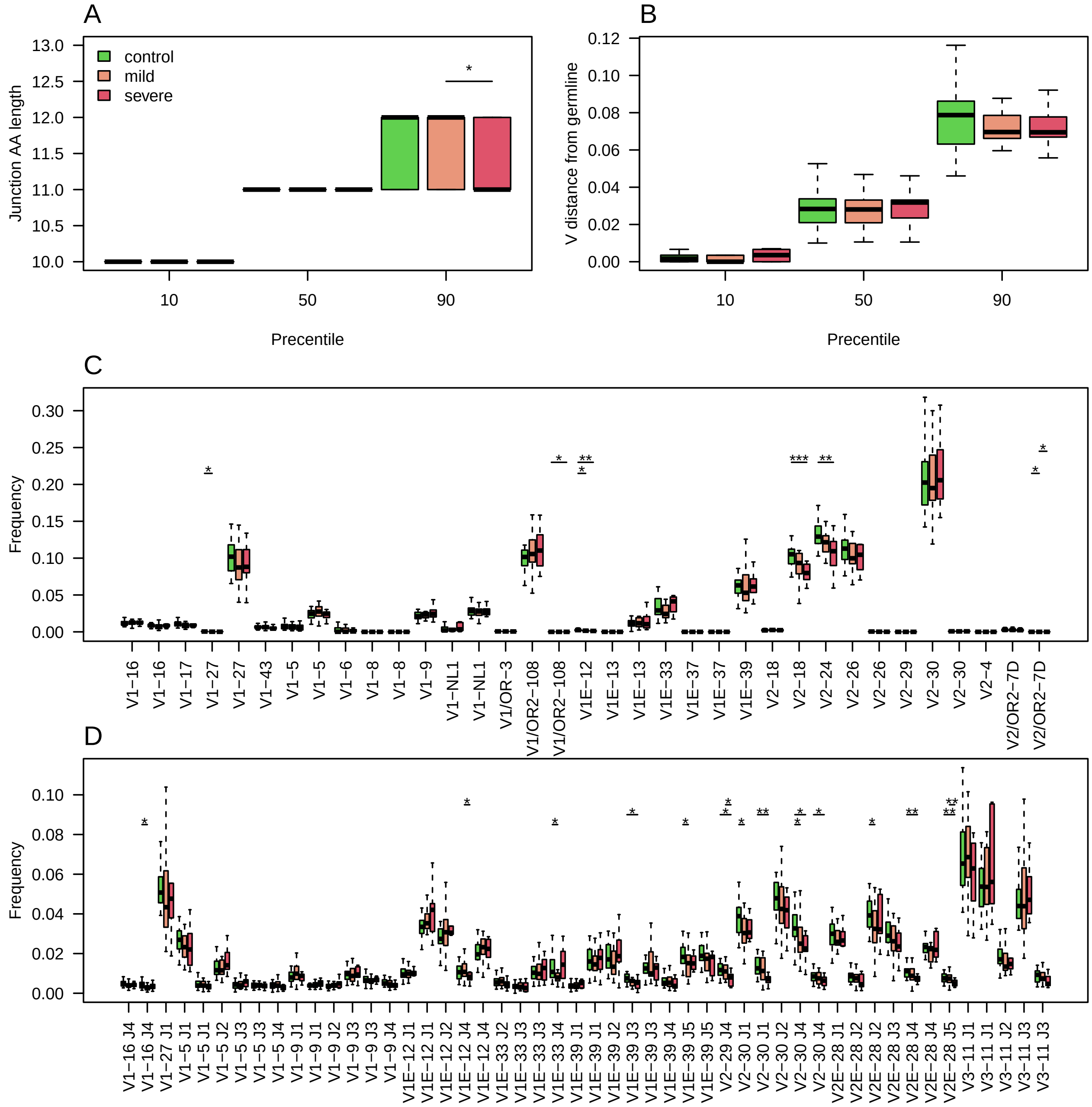


Figure S2: **Characteristics of the light kappa chain sequencing data**
A. 10,50 and 90 percentiles of AA CDR3 lengths in individuals with COVID-19 at indicated severity levels and healthy controls. B. 10,50 and 90 percentiles of V gene distances from germline in individuals with COVID-19 at indicated severity levels and healthy controls. C. Boxplots showing V gene usage in individuals with COVID-19 at indicated severity levels and healthy controls. Shown are the top 50 highest mean frequencies. D. V & J gene usage comparison. Throughout the figure, * marks a P value lower than 0.05, ** marks a P value lower than 0.01, and *** marks a P value lower than 0.001.

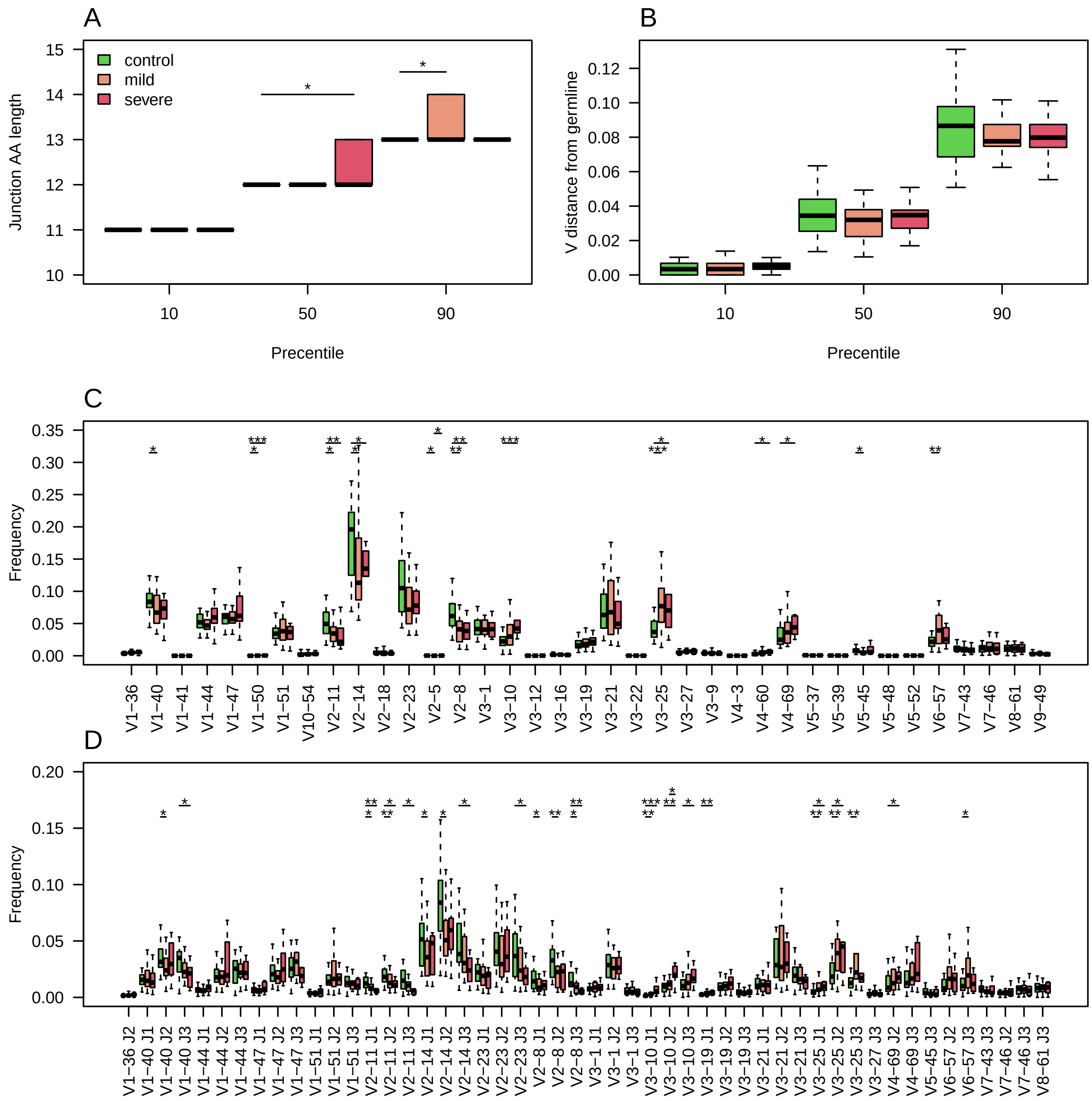


Figure S3: **Characteristics of the light lambda chain sequencing data**
A. 10,50 and 90 percentiles of AA CDR3 lengths in individuals with COVID-19 at indicated severity levels and healthy controls. B. 10,50 and 90 percentiles of V gene distances from germline. C. Boxplots showing V gene usage. Shown are the top 50 highest mean frequencies. D. V & J gene usage comparison. Shown are the top 50 highest mean frequencies. Throughout the figure, * marks a P value lower than 0.05, ** marks a P value lower than 0.01, and *** marks a P value lower than 0.001.

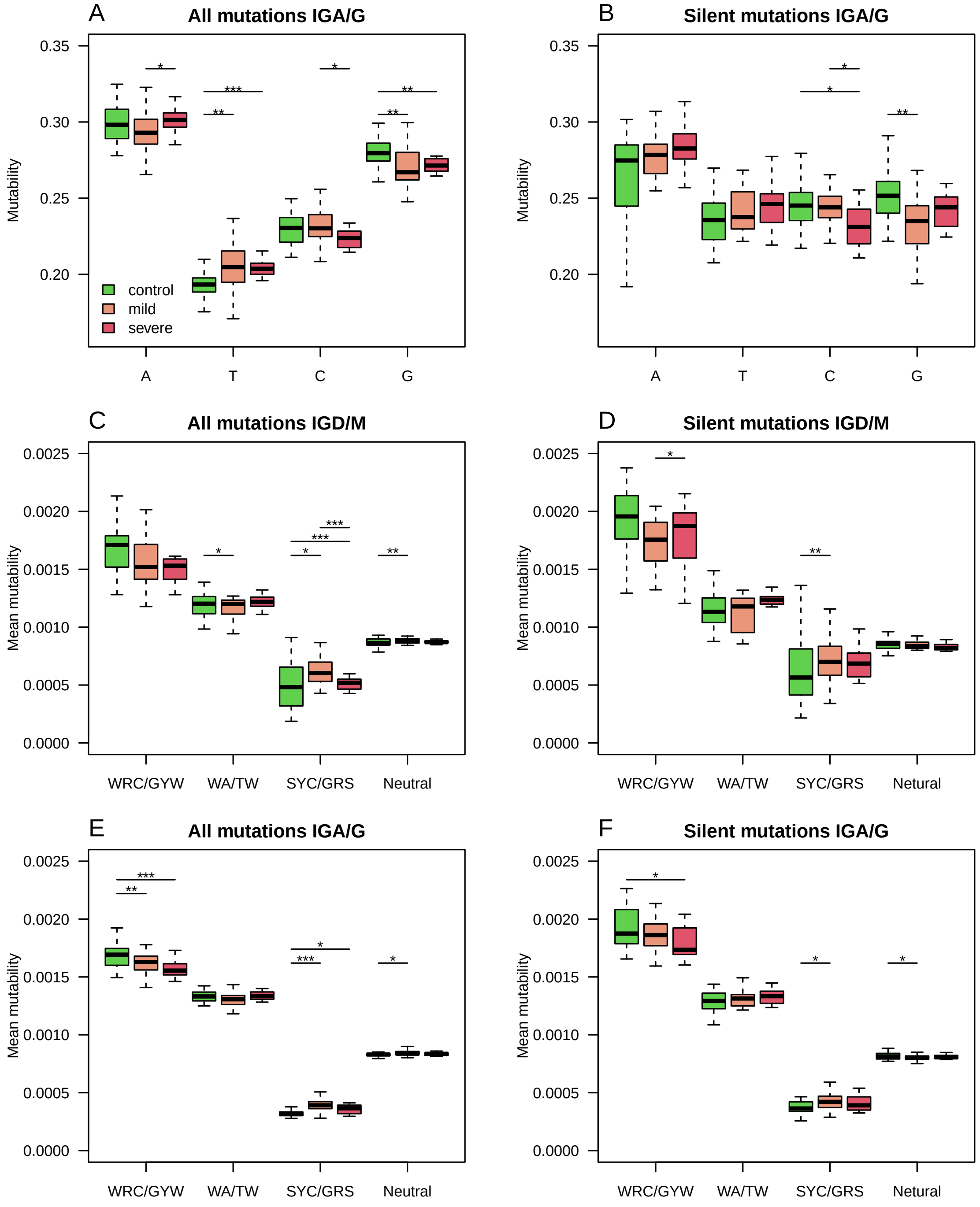


Figure S4: **Silent and replacement mutability in SHM: single base mutability, 5-mers hot-spots and cold-spots**
A. A single base mutability model was built based on IGA/G isotypes of COVID-19 patients and controls, taking into account only one representative from each clone. Shown are boxplots representing the normalized sum of single base mutability. B. The same plot as in A, but for silent mutations only. C-D. A 5-mer SHM model based on both silent and replacement mutations in C, or silent only mutations in D, was built using the IGD and IGM isotypes of COVID-19 patients at different severity levels and healthy controls. Shown are the known SHM hot-spots, SHM cold-spots, and the rest of the sites. E-F. A 5-mer SHM model based on both silent and replacement mutations in E, or silent only mutations in F, was built using the IGA and IGG isotypes of COVID-19 patients at different severity levels and healthy controls. Shown are the known SHM hot-spots, SHM cold-spots, and the rest of the sites. Throughout the figure, * marks a P value lower than 0.05, ** marks a P value lower than 0.01, and *** marks a P value lower than 0.001.

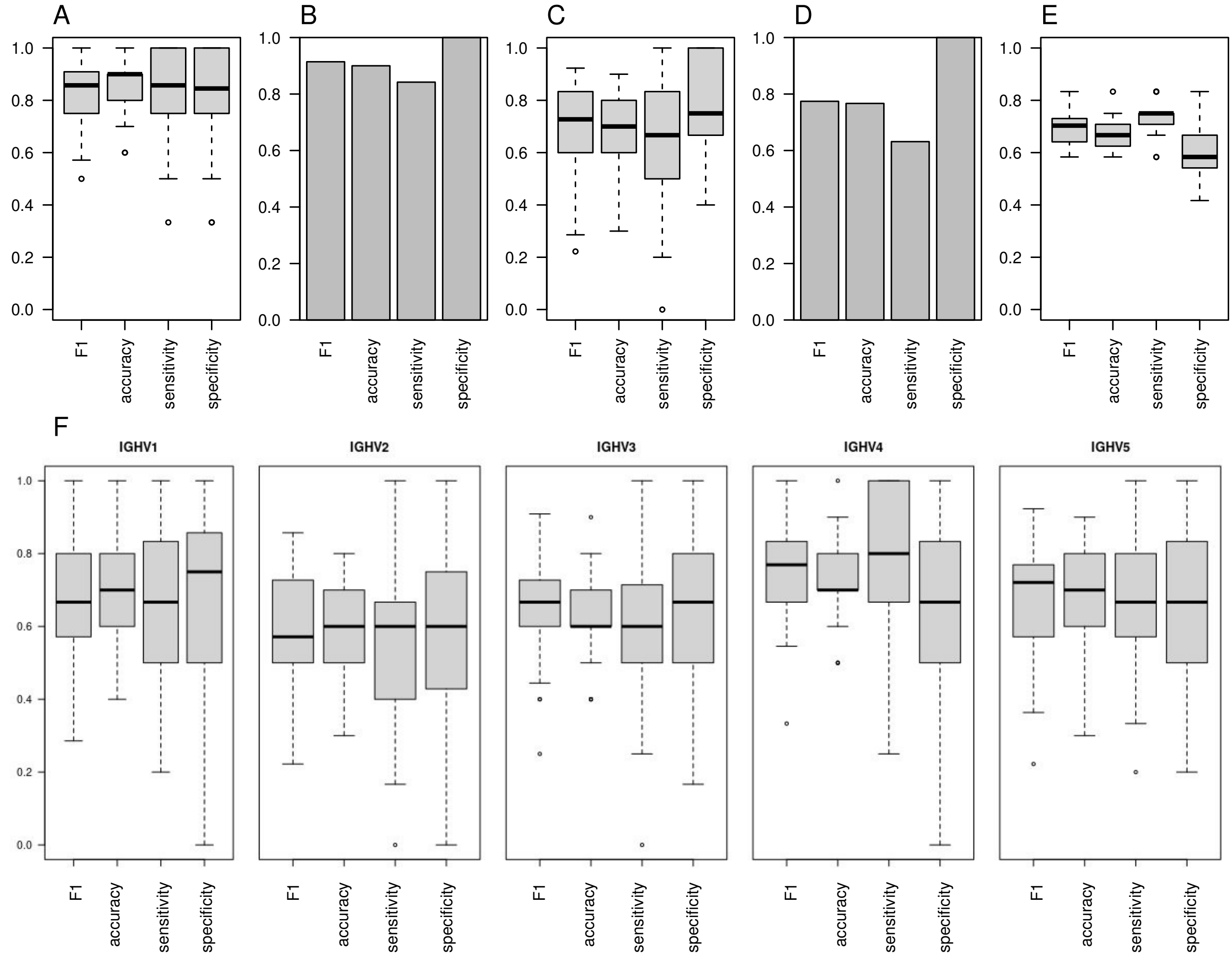


Figure S5: **SHM in the heavy chain enables both COVID-19 classification and severity classification using one representative from each clone, but to less efficiency when building the matrix based on a single V family**

A. An ML algorithm was trained on the substitutions matrix of the 5-mer SHM model (taking into account only one representative from each clone), which was created for the IGA/G isotypes. Boxplots representing F1 score, accuracy, specificity, and sensitivity of 50 random splits to train and test groups are shown. B. Logistic regression was trained on the substitutions matrix of the SHM model built from the entire dataset. Barplot representing F1 score, accuracy, specificity, and sensitivity of classifications on the test group. C. The same algorithm as in A was trained on silent mutations only. Shown are Boxplots representing the F1 score, accuracy, specificity, and sensitivity of 50 random splits to train and test groups. D. The same algorithm as in C was trained on silent mutations only. Shown are barplots representing the F1 score, accuracy, specificity, and sensitivity of classifications on the test group. E. Boxplots showing F1 score, accuracy, specificity, and sensitivity of 20 leave-one-out cross validation of severity classification. Each leave-one-out was on 12 severe COVID-19 patients and 12 randomly selected mild COVID-19 patients. The ML algorithm was trained on the mutability matrix of the SHM cold-spots in these groups. F. F1 score, accuracy, specificity, and sensitivity of classifications based on single V family SHM matrices.

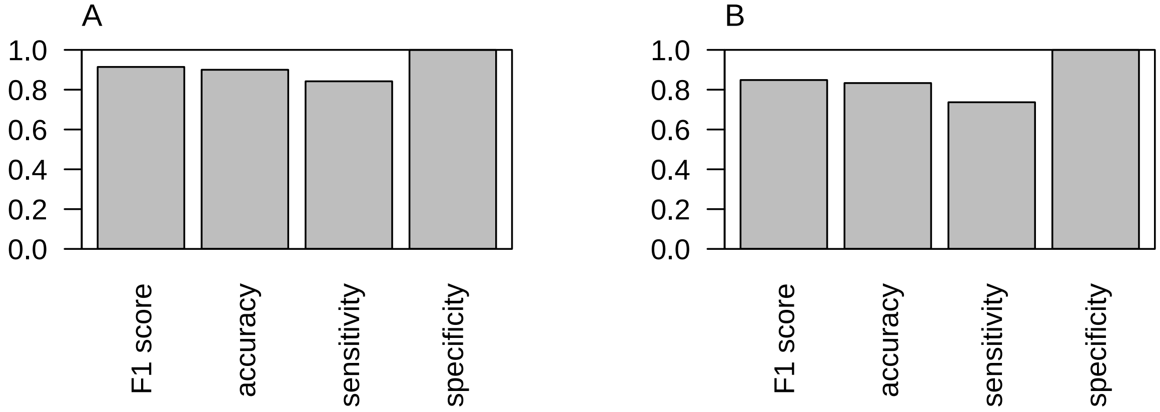


Figure S6: **SHM of heavy chains enables COVID-19 classification - test group**
A. Logistic regression was trained on the substitutions matrix of SHM model built from the entire dataset. A barplot representing F1 score, accuracy, specificity, and sensitivity of classifications on the test group. B. The same algorithm as in A was trained on silent mutations only. Shown are barplots representing the F1 score, accuracy, specificity, and sensitivity of classifications on the test group.

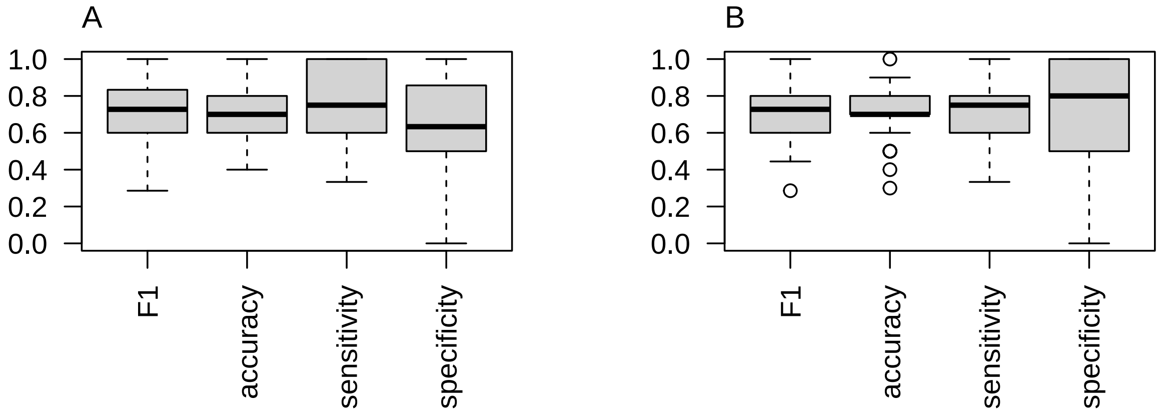


Figure S7: **SHM of light chains enables COVID-19 classification**
A. Logistic regression was trained on the substitutions of SHM model built from the entire dataset. Boxplot representing F1 score, accuracy, specificity, and sensitivity of 50 random splits to train and test groups are shown. B. The same algorithm as in A was trained on silent mutation only. Shown are boxplots representing the F1 score, accuracy, specificity, and sensitivity of 50 random splits to train and validation groups.

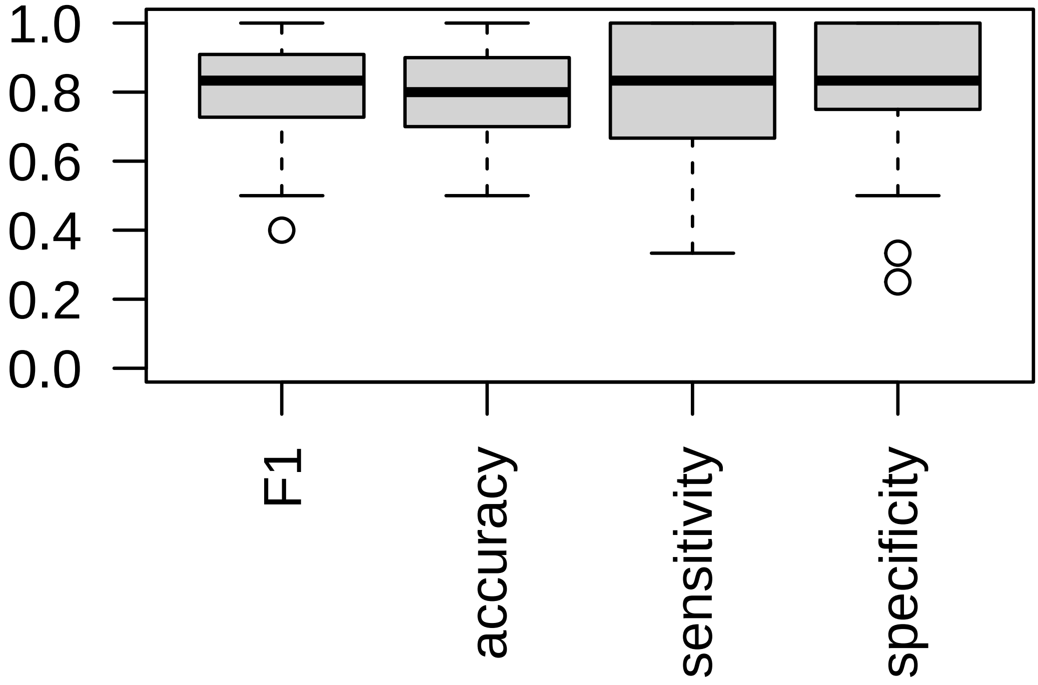


Figure S8: **SHM both Heavy and Light chains enables COVID-19 classification** A. Logistic regression was train on the substitutions of SHM model built on data. A boxplot representing F1 score, accuracy, specificity, and sensitivity of classifications of 50 random splits to train and validation groups are shown.

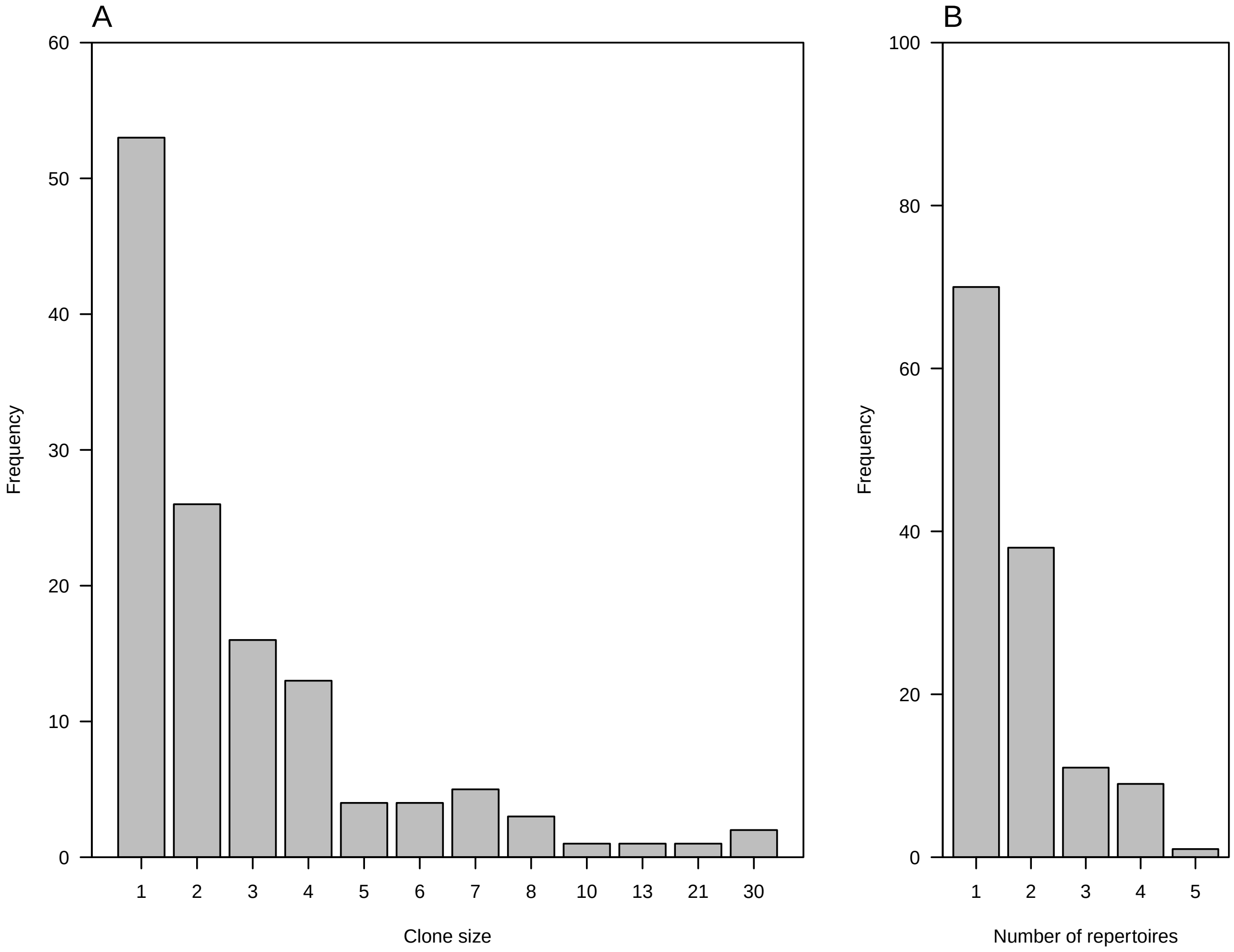


Figure S9: **Characterization of known clones of COVID-19 antibodies**
A. The frequencies of clones found in our COVID-19 patients with indicated clones sizes. B. The frequencies of clones found in our COVID-19 patients with the indicated number of repertoires having at least one sequence which belongs to the clone.

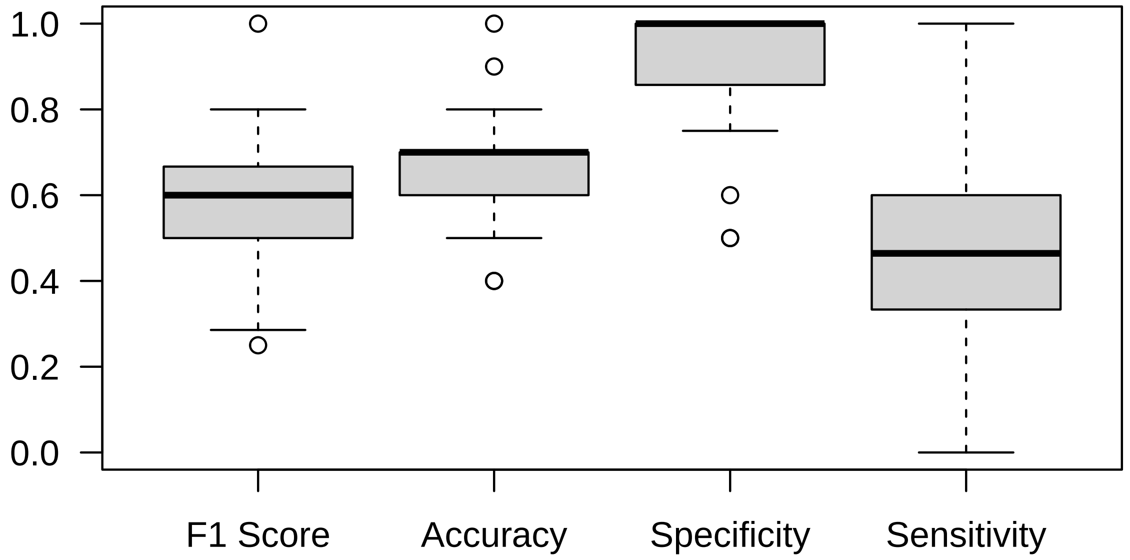


Figure S10: **B cells shared clones enable weak COVID-19 classification** Samples were randomly split to train and validation groups. Shared clones were counted in the training group, and logistic regression was trained on tables summarizing the frequency of each clone in all training samples. Classifications were then made for the validation group. A Boxplot representing the F1 score, accuracy, specificity, and sensitivity of classifications of 50 random splits to train and validation groups is shown.
